# PERK inhibition with GSK2606414 in zebrafish evokes developmental defects consistent with Wolcott-Rallison syndrome phenotypes

**DOI:** 10.1101/2024.04.16.589737

**Authors:** Liliana M. Almeida, Leonor Pereira Lima, Nuno A. S. Oliveira, Rui F. O. Silva, Bruno Sousa, José Bessa, Brígida R. Pinho, Jorge M. A. Oliveira

## Abstract

**Background:** PERK (EIF2AK3) is an endoplasmic reticulum stress kinase whose loss of function disturbs human development, leading to skeletal dysplasia and permanent neonatal diabetes, as observed in the Wolcott-Rallison Syndrome (WRS). The lack of effective, less invasive therapies for developmental diseases highlights the need for animal models that replicate complex pathological phenotypes, while allowing scalable drug screening. Zebrafish, with their high fecundity and rapid development, facilitate efficient *in vivo* drug testing.

**Methods:** We aimed to assess the potential of zebrafish for studying PERK function and its pharmacological modulation, particularly as a model for developmental diseases like WRS. Bioinformatic analyses assessed the similarity between human and zebrafish PERK. Increasing concentrations of GSK2606414 were used to inhibit PERK. A combination of behavioural and functional assays evaluated the effects of GSK2606414 on zebrafish skeletal, neuromuscular, and cardiac development. Fluorescence microscopy in transgenic zebrafish expressing fluorescent pancreatic markers and a glucose probe assessed the diabetic-like phenotype.

**Results:** We found high similarity between human and zebrafish PERK, along with bioactivity of the PERK inhibitor GSK2606414 in zebrafish. PERK inhibition evoked defects in WRS relevant parameters, such as growth and skeletal development, as well as neuromuscular and cardiac deficiencies, whereas parameters not associated with WRS like otolith area and eye/body ratio remained unaffected. Moreover, PERK inhibition decreased pancreatic ! cell mass and disrupted glucose homeostasis, indicating a diabetic phenotype.

**Conclusion:** These findings evidence zebrafish’s potential for studying PERK function and its pharmacological modulation in developmental disorders like WRS, aiding research on pathophysiology and experimental treatments.

## 1. Introduction

Protein kinase RNA-like endoplasmic reticulum kinase (PERK) is an endoplasmic reticulum (ER) membrane kinase that activates in response to disruptions in ER protein homeostasis (proteostasis) by promoting a protective response, the integrated stress response (ISR). Upon activation by stress, PERK phosphorylates eukaryotic translation initiation factor 2α (eIF2α) that decreases global protein synthesis, alleviating nascent protein load (1) and boosts the synthesis of transcription factors like activating transcription factor 4 (ATF4) and C/EBP homologous protein (CHOP). These transcription factors regulate the expression of chaperones, proteases and components of the autophagic pathway, thus contributing to proteostasis maintenance (1). During development, protein synthesis plays a critical role in cell division, differentiation, and the rapid growth of embryos (2), and PERK is essential for detecting protein misfolding and adjusting protein synthesis to maintain proteostasis (3).

In humans, loss-of-function mutations in the *EIF2AK3* gene encoding for PERK cause a rare (prevalence of <1/1 000 000; ORPHA:1667) autosomal recessive developmental disease named Wolcott-Rallison Syndrome (WRS), with patients dying before 35 years of age (4). WRS is mainly characterized by skeletal dysplasia and neonatal diabetes mellitus (5), but WRS patients may also present broad symptomatology like motor neuropathy, intellectual deficit, early neurodegeneration, and cardiac malformations (4). This broad symptomatology is likely due to the widespread distribution of PERK, with several loss-of-function mutations totally or partially abolishing its kinase activity (4, 6). Existing therapies for WRS patients include blood glucose control with insulin pumping and transplant of liver/pancreas and kidney (7). The scarcity of non-invasive and more effective therapeutic options for WRS patients highlights the need for further therapeutic investigation, which requires efficient WRS models.

The currently available *in vivo* models for the developmental disease WRS are rodents with PERK knock-out or loss-of-function mutations in PERK that exhibit the WRS hallmarks– neonatal diabetes and skeletal dysplasia (8, 9) – and the broad-spectrum phenotypes – cardiac, muscle and cognitive abnormalities (10–12). However, for large exploratory studies of new therapeutics, there are efficiency advantages in using small vertebrates like zebrafish before testing the most promising candidates in rodent models. Zebrafish are increasingly used to develop disease models given their advantageous features: high fecundity, external fertilization, and rapid formation of complete organs, making zebrafish ideal for investigating developmental diseases and for rapid and scalable *in vivo* drug testing (13, 14). Previous studies that targeted PERK in zebrafish, used either genetic (morpholino knockdown) or pharmacological inhibition (15–17). In such studies, PERK inhibition with GSK2606414 was used in zebrafish adults to study sleep regulation (15), and PERK inhibition with GSK2656157 or PERK knockdown was used to study redox regulation and neurogenesis in zebrafish embryos (16, 17). To the best of our knowledge, there are no previous studies in zebrafish with a primary focus on studying PERK function in developmental diseases, including WRS.

Here, we combined bioinformatic analyses, molecular biology techniques, behaviour, and functional assays to assess the effects of pharmacological PERK inhibition in zebrafish. We used transgenic zebrafish expressing pancreatic reporters and a fluorescent glucose probe to assess diabetic-like phenotypes. We performed functional and structural microscopy to assess skeletal, neuromuscular, and cardiac development. Our data show that zebrafish holds potential as a model to study PERK function in developmental disorders, such as WRS.

## 2. Material and Methods

### 2.1 Drugs, solvents, and solutions

Stock solutions of the PERK inhibitor GSK2606414 (Merck Millipore, 516535) (15, 18), the muscle relaxant D-tubocurarine (dTC; Sigma, T2379), and the anesthetic tricaine methanesulfonate (0.02%; Sigma, E10521) were prepared in DMSO (Sigma, 276855) and diluted in standard E3 medium (19). DMSO levels were kept under 0.2% (20).

### 2.2 Zebrafish maintenance and husbandry

Adult AB strain zebrafish were supplied by the Interdisciplinary Centre of Marine and Environmental Research (CIIMAR, Portugal), transgenic strains of zebrafish were obtained from José Bessa’s laboratory at the Institute for Research and Innovation in Health (i3s, University of Porto, Portugal). Zebrafish adults were maintained at 28 ± 1°C on a 14 h:10 h light:dark cycle (21, 22). Breeding pairs (1:1, male:female) were placed in breeding tanks on the day before egg collection. Ninety minutes after starting the light period, eggs were collected, and time point was recorded as 0 h post fertilization (hpf). **Ethic statement:** Handling of zebrafish followed the European Directive 2010/63/EU and the Portuguese Law (Decreto-Lei 113/2013). Experiments were performed according to the 3Rs principle by reducing the number of animals used and using refined techniques that reduce animal suffering, using procedures approved by the i3S Animal Welfare Body.

### 2.3 Drug treatments

5 zebrafish/well were randomly distributed in 24-well plates (0.5 mL E3/well) and kept at 28 ± 1°C on a 14 h:10 h light: dark cycle throughout the assay. Zebrafish were continuously exposed to GSK2606414 or solvent (DMSO) from 4 to 76 hpf and solutions were renewed daily (21). dTC was incubated for 10 minutes before assays endpoint (≥ 15 zebrafish).

### 2.4 Brightfield microscopy

For brightfield microscopy, we used a stereomicroscope (Stemi DV4, Zeiss, Göttingen, Germany) or an inverted microscope (Eclipse TE300, Nikon, Tokyo, Japan). The inverted microscope was attached to a motorized stage (ProScan, Prior) and a CCD camera (ORCA-ER, Hamamatsu), constituting a Nikon-Prior-Hamamatsu (NPH) imaging system that was controlled by the open-source software Micro-Manager (v. 2.0; https://micro-manager.org):

#### 2.4.1 Monitoring zebrafish development

Zebrafish were scored as normal, abnormal, or dead; hatched (ruptured chorion and free tail movement) or unhatched at 28, 52 and 76 hpf. Death was ascertained according to OECD guideline 236 (23), normal development as previously described (24), and abnormalities as cardiac and skeletal (further into scoliosis, curly-up tail, or kyphosis). Dead zebrafish were removed at each time point and only live ones were used in subsequent assays.

#### 2.4.2 Morphometric analysis

Zebrafish body length was measured from the tip of the head (between the eyes) to the end of the tail (25) and trunk-tail angle was measured at the intersection between the rostral (trunk) and caudal (tail) notochord at the level of the anus (Figure 3 B).

#### 2.4.3 Heartbeat and atrio-ventricular coordination

Heartbeats were counted in 4 zebrafish per condition, per experiment, during 20 s, at 72-80 hpf (26). Atrio-ventricular coordination recordings were obtained with “Plot Z-axis profile” on ImageJ after imaging zebrafish for 90 seconds at 12 frames per second.

#### 2.4.4 Tail coilings

Spontaneous activity was measured at 28 hpf in glass bottom 96-well plates where each zebrafish was imaged for 3 minutes at a frequency of 4.5 frames per second and 2×2 or 4×4 binning (19).

#### 2.4.5 Birefringence analysis

76 hpf zebrafish were anesthetized, immobilized on a glass surface, oriented laterally, and illuminated with linearly polarized light (Screen-Tech, ST-38). An orthogonal polarization filter was placed after the samples to block directly transmitted light. Thus, only light whose polarization angle is affected by the sample is collected by the NPH imaging system (27). Pixel intensity in the trunk region was measured by selecting the region of interest with ImageJ software.

### 2.5 Touch-evoked escape response

At 76 hpf, at least 6 zebrafish/condition were submitted to 10 alternate touches to the head and to the tail with 10 µL pipette tip with a recovery time of 30 seconds (28). Data are presented as response rates in percentage of total touches.

### 2.6 Fluorescent microscopy

Zebrafish were imaged with the NPH system coupled to a monochromator (Polychrome II, Photonics) as follows.

#### 2.6.1 2-NBDG labelling

3 hours before 76 hpf end-point, zebrafish were incubated with a fluorescent glucose analogue - 600 µM 2-NBDglucose (2-(7-Nitro-2,1,3-benzoxadiazol-4-yl)-D-glucosamine) (Tebu-Bio,10-1459) (29, 30); and a positive control for neuromast labelling - Mitotracker Deep Red (1 µM, Invitrogen, M22426, labels polarized mitochondria in neuromasts) in 96 well plates (4 zebrafish /well). At 76 hpf, each zebrafish was anesthetized and imaged in a glass-bottom chamber. 2-NBDglucose was excited at 475 nm and Mitotracker Deep Red at 650 nm for 1 second.

Emissions were collected using band pass filters (Chroma 69008 – ET – ECFP/EYFP/mCherry) with a 10 × objective. Images of the entire fish were stitched with the GNU image manipulation program (The GIMP Development Team, 2019; https://www.gimp.org), before evaluation of anterior and posterior neuromasts (31).

#### 2.6.2 Transgenic zebrafish

Double transgenic zebrafish expressing GFP in insulin-producing β cells and mCherry in somatostatin-producing δ cells (Tg(ins:GFP, sst:mCherry)) or in elastase-producing cells (Tg(ins:GFP, ela:mCherry)) were kept in 24-well plates (5 zebrafish/plate) with E3 media. We used two pharmacological approaches: 1) GSK2606414 exposure from 4 hpf (before the principal islet of Langerhans is formed within the endocrine pancreas) to 28, 76 or 148 hpf; 2) GSK2606414 exposure from 76 hpf (when the principal islet is already formed) to 148 hpf (32). At the final time point (28, 76 or 148 hpf) zebrafish were anesthetized by rapid cooling on ice and fixed in 4% paraformaldehyde overnight. GFP was excited at 488 nm and mCherry was excited at 575 nm. Emissions were collected using band pass filters (Chroma 59022 – EGFP/mCherry) with a 20 × objective. The area of GFP- or mCherry-labelled pancreas in each image was assessed with the ImageJ software (https://imagej.nih.gov/ij).

### 2.8 Western blot

24-27 zebrafish larvae were rinsed with ice-cold water twice before being placed in RIPA buffer (50 mM Tris (NZYTech, MB01601), 150 mM NaCl (Merck, S9888), 1 mM EDTA (Sigma-Aldrich, E6758), 1% IGEPAL CA-630 (Sigma-Aldrich, I3021), 1% sodium deoxycholate (Sigma-Aldrich, D6750), 0.1% sodium dodecyl sulfate (Sigma-Aldrich, L3771) supplemented with a protease and phosphatase inhibitor cocktail (Thermo Scientific, 78440). After 3 freeze-thaw cycles, homogenization with Precellys Evolution (6800 rpm, 5 × 42 s with 37 s interval) and centrifugation at 12,000 × g (10 min, 4 °C), supernatant protein concentrations were quantified by the Bradford method. 30 to 40 µg of protein extracts were supplemented with Bolt LDS sample buffer (Invitrogen, B0007) and Bolt sample reducing agent (Invitrogen, B0009), denatured at 95 °C for 5 min, and loaded into 4–15% Mini-PROTEAN TGX™ Precast Protein Gels (Biorad, 4561084), which were electrophoresed (125 and 200 V, 30 min) and wet-transferred to PVDF membranes (Millipore, IPVH00010) overnight (30 V, 4°C). Membranes were blocked (1 h, room temperature or overnight at 4°C) with 5% bovine serum albumin (BSA, NZYTech, MB04602) in PBS with 0.05% Tween 20 (PBST) and incubated with primary antibodies (Supplementary Table 1). After washing with PBST, membranes were incubated with respective horseradish peroxidase-conjugated secondary antibodies (1 h) (Supplementary Table 2), for detection with a Chemiluminescent kit (Novex ECL, Invitrogen, WP20005) and a ChemiDoc MP Imaging system (Bio-Rad). Membrane Coomassie (Sigma-Aldrich, 27816) staining was used for loading control rather than ‘housekeeping’ proteins, as their levels may change under certain conditions (33). Densitometric analyses were performed with the Image J software. In each independent experiment, samples from control (0 µM GSK) and 10 µM GSK-treated zebrafish were run in the same gel.

### 2.9 Bioinformatic analyses

#### 2.9.1 Comparison of PERK orthologues

Protein sequence data from zebrafish (*Danio rerio,* XP_005156642.2), human (*Homo sapiens,* AAI26357.1), mouse (*Mus musculus,* AAH54809.1), and fruit fly (*Drosophila melanogaster,* AAF61200.1) were obtained from UniProt (https://www.uniprot.org/, 28^th^ February 2025). The amino acid sequences from the total human PERK protein and from its kinase domain were compared with its orthologue proteins from zebrafish, mouse, and fruit fly, using the Protein BLAST tool (https://blast.ncbi.nlm.nih.gov/Blast.cgi). Additionally, we determined the identity of the vital amino acids for GSK2606414 binding to the catalytic site of PERK, by aligning the protein sequence from zebrafish and human.

#### 2.9.2 Zebrafish PERK 3D structure and molecular docking

The predicted three-dimensional protein structures of zebrafish PERK (UniProt: B8JI14) and human PERK (UniProt: Q9NZJ5) were obtained from AlphaFold Protein Structure Database (34, 35). UCSF ChimeraX software was used to visualize, align (Matchmaker tool) and calculate the root-mean-square deviation (RMSD) of the three-dimensional structures of human PERK (PDB: 4g31) and zebrafish PERK (UniProt: B8JI14), and GSK2606414.

### 2.10 Statistical Analyses

Data are shown as mean ± standard error of the mean (SEM) of the *n* specified in figure legends. For experiments with only two groups, comparisons were made with Student’s *t*-test. For single-factor (drug concentration) comparisons of multiple groups to their respective control, we used One-Way ANOVA with Dunnett’s *post-hoc* test. For two-factor (drug concentration vs. time) analysis, we used Two-Way ANOVA with Sidak’s *post-hoc* test. For survival analysis we used the Mantel-Cox (Log-rank) test. *P* < 0.05 was taken as statistically significant. Data analyses were performed with Prism 6.0 (GraphPad).

## 3. Results

### 3.1 The PERK-GSK2606414 binding site is conserved in humans and zebrafish

To assess the potential of zebrafish to study PERK function and its pharmacological modulation, we performed bioinformatic comparisons of the amino acid and structural identity of zebrafish and human PERK, including specifically the binding site for the PERK inhibitor GSK2606414. We selected GSK2606414 (hereafter GSK; not to be confused with glycogen synthase kinase) because: it is one of the most used PERK inhibitors (1); it was previously used in zebrafish (15); and the vital amino acids for its binding to PERK are identified (36, 37).

The zebrafish PERK homologue presented 59% global amino acid identity with human PERK (Figure 1 A). Using Protein BLAST, we observed that out of the 12 vital amino acids for GSK binding to PERK (36, 37), there is only one difference between human and zebrafish (Figure 1 B). Such difference is a valine in human vs. isoleucine in zebrafish, and both amino acids are hydrophobic. Using molecular visualization with ChimeraX, we observed that the average distance between the 3D structures of zebrafish and human PERK is under 1 Å (Figure 1 C, D), indicating high similarity (38).

**Figure 1.**
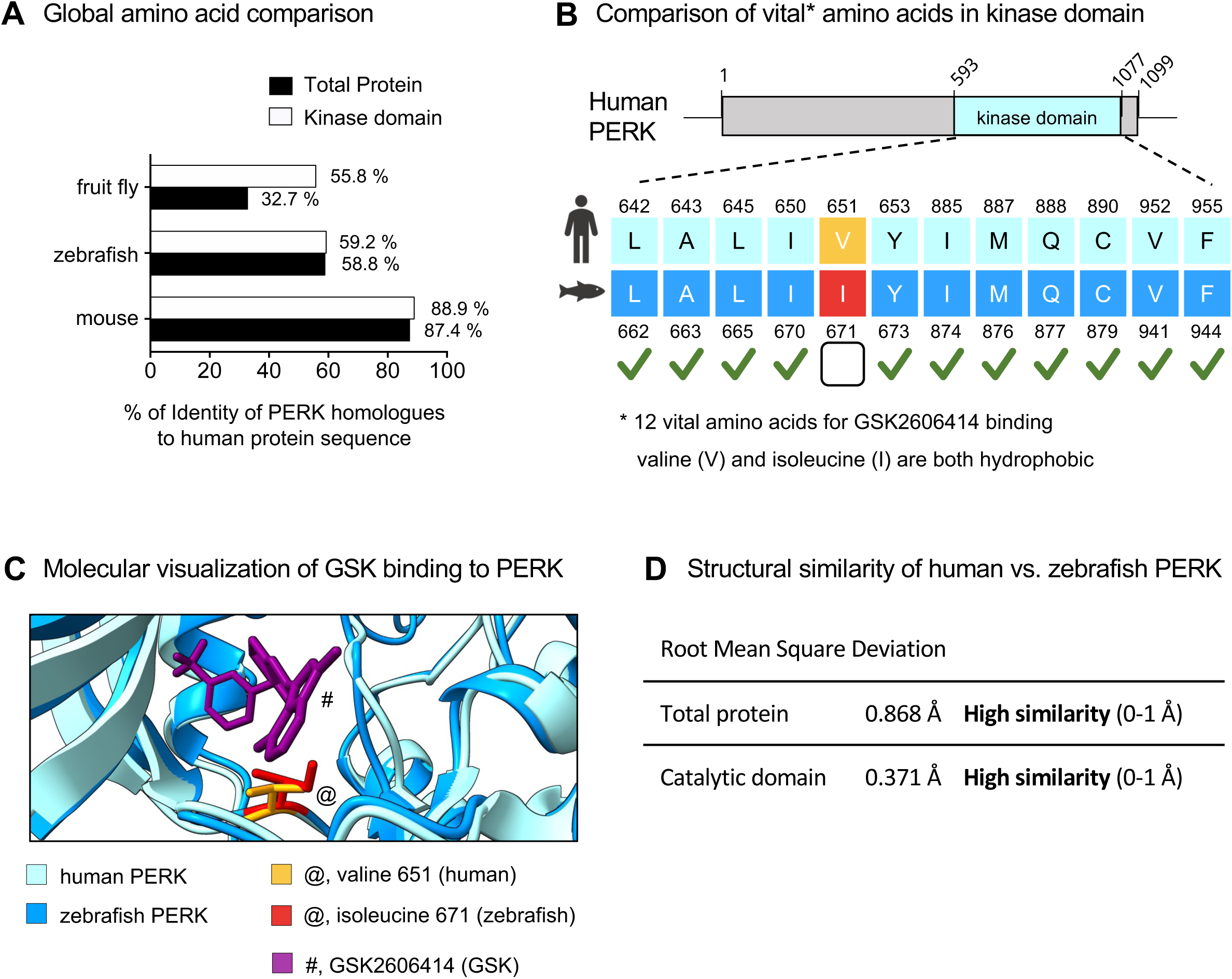
Bioinformatic analyses of PERK identity and GSK2606414-binding in human vs. zebrafish. (**A**) Percentage of identity between human, fruit fly (*Drosophila melanogaster*), zebrafish (*Danio rerio*), and mouse (*Mus musculus*) PERK (total and kinase domain); Protein BLAST. (**B**) Comparing vital GSK2606414 (GSK)-binding amino acids in the catalytic domain of human and zebrafish PERK. Valine 651 in human (yellow) is substituted by the also hydrophobic Isoleucine 671 (red) in zebrafish; Protein BLAST. (**C**) Predicted GSK (purple, #) binding to human (light blue) and zebrafish (dark blue) PERK; ChimeraX. (**D**) Root Mean Square Deviation (RMSD) comparing human and zebrafish PERK tertiary structures and respective reference values; ChimeraX.

These results predict a high probability of GSK binding and inhibiting zebrafish PERK. To test such prediction, we proceeded with functional assays.

### 3.2 The PERK inhibitor GSK2606414 is bioactive in zebrafish

To evaluate the effects of PERK inhibition, we evaluated the survival and hatching of zebrafish treated with increasing concentrations of GSK over time (Figure 2 A). We observed a time-dependent decrease in survival, with 10 μM GSK being the concentration that induces a significant effect (50%) at 76 hpf (Figure 2 B). For zebrafish that survived GSK, hatching increased time-dependently, without significant differences across GSK concentrations (Figure 2 C). Next, we assessed the effect of GSK on the levels of PERK pathway (ISR) biomarkers in the zebrafish at 76 hpf using western blot (Figure 2 D *i*). Treatment with GSK (10 μM, 72h) significantly decreased levels of phosphorylated eIF2α, ATF4, and CHOP (Figure 2 D *ii-vi*). Together, GSK effects confirm its bioactivity and are compatible with PERK pathway inhibition as previously reported (15).

**Figure 2.**
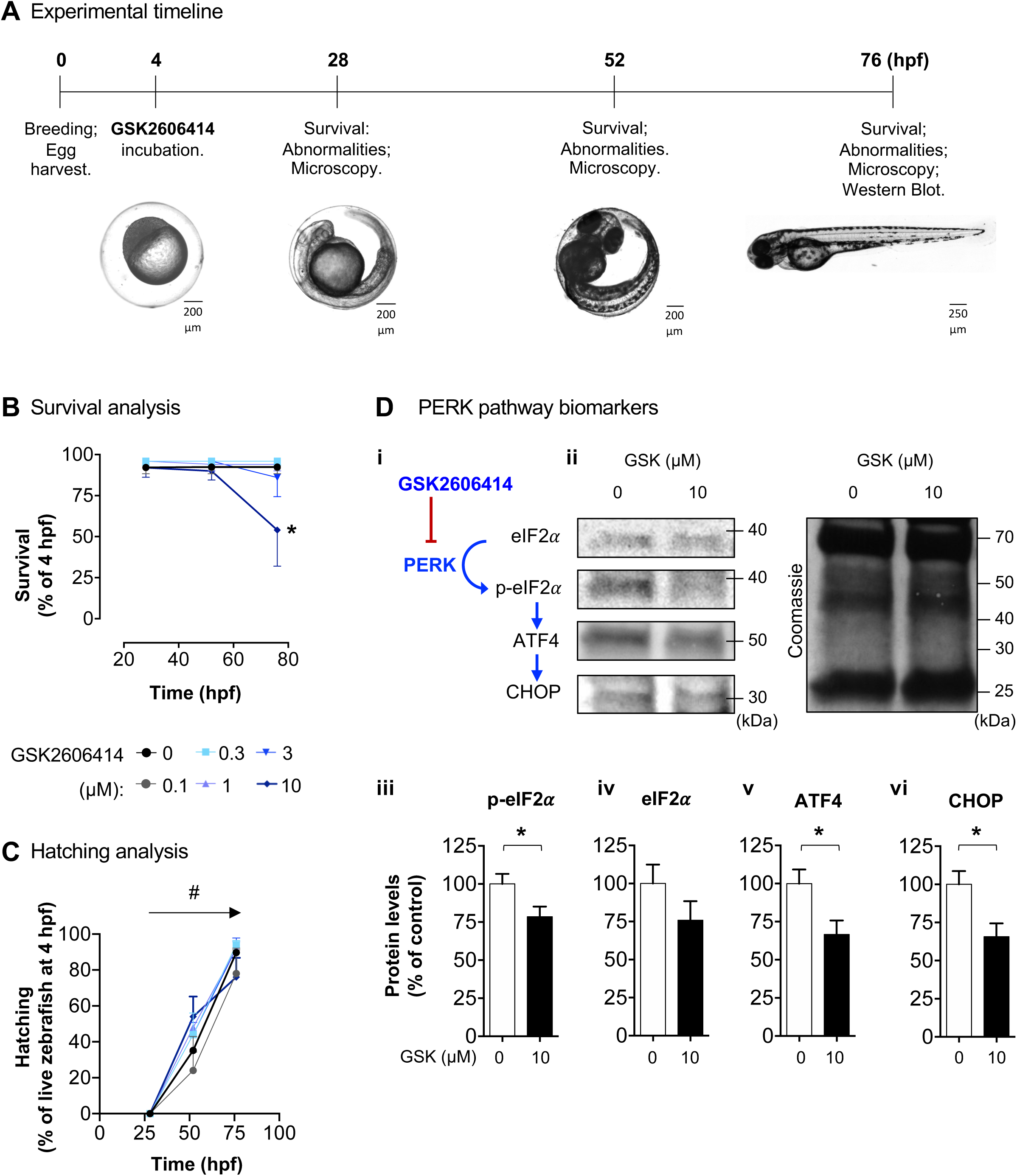
GSK2606414 effects on zebrafish survival, hatching, and PERK pathway biomarkers. (**A**) Experimental timeline. (**B**) Zebrafish survival when exposed to GSK2606414 (GSK; 0-10 μM) over time (4-76 hours post-fertilization; hpf); *n* = 50-234 zebrafish, 5 independent breedings, Mantel-Cox test for survival, \**P* < 0.05 vs. 0 μM GSK (control). (**C**) Hatching of zebrafish treated with GSK; *n*= 50-234 zebrafish, 5 breedings, Two-Way ANOVA, time effect: ^#^*P* < 0.05. (**D**) Western blot of live 76 hpf zebrafish: *i*) Schematic representation of the PERK pathway biomarkers; *ii-vi*) Representative blots and quantifications; *n* = 4 independent experiments (pool of 24-27 larvae per condition and per experiment); *t*-test: \**P* < 0.05.

**Figure 3.**
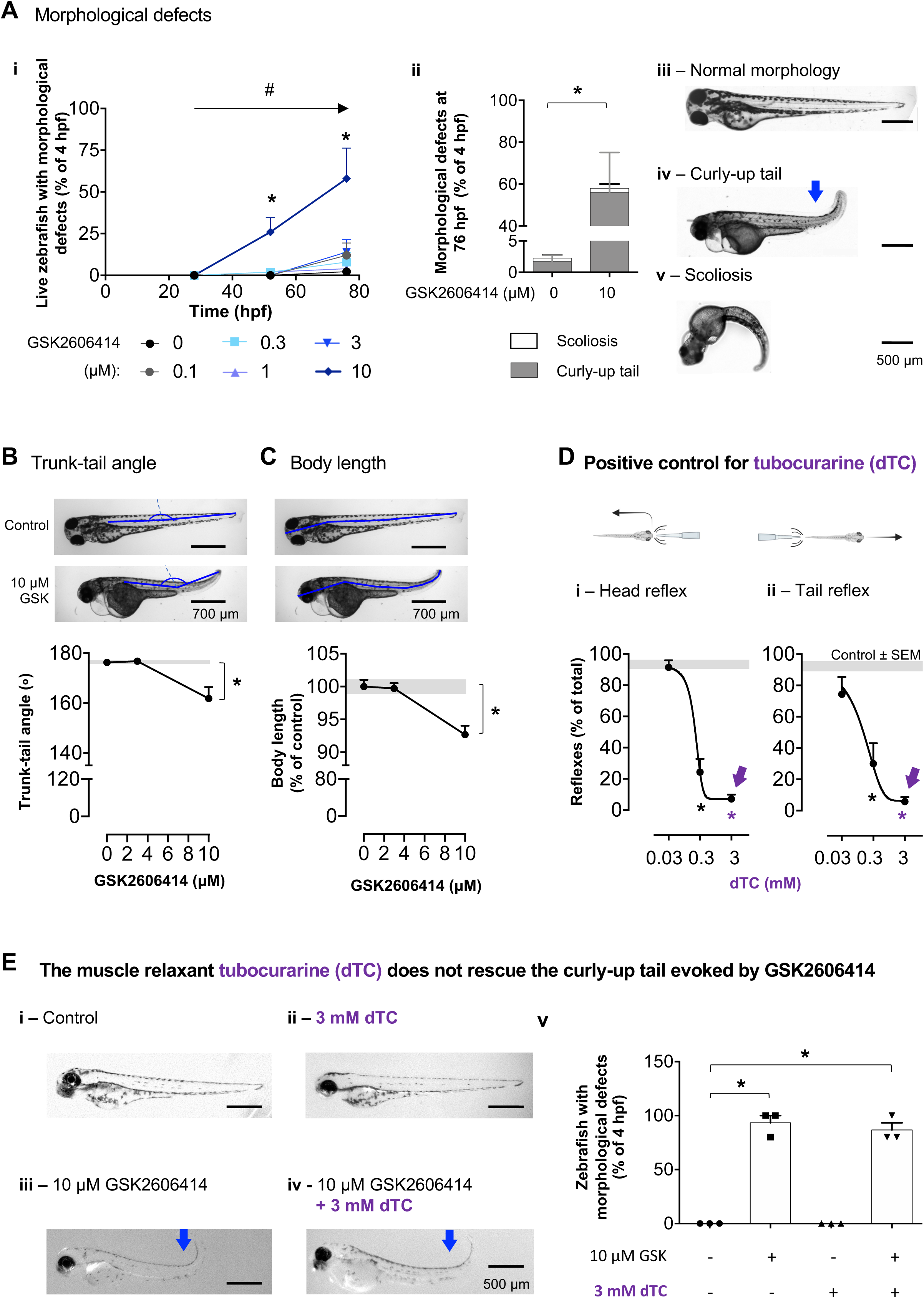
GSK2606414 impaired skeletal development in zebrafish. (**A**) (*i*) Morphological defects following GSK2606414 (GSK) (0-10 μM) treatment overtime; \**P <* 0.05, 10 vs. 0 µM GSK; ^#^*P<*0.05, effect of time for 10 µM GSK, Two-Way ANOVA with Sidak *post-hoc*; *n* = 50-234 zebrafish; 5 independent experiments. (*ii-v*) Proportion of curly-up tail and scoliosis at 76 hpf, and representative images. (**B, C**) Measurements (blue lines) and quantifications of trunk-tail angle and body length at 76 hpf; \**P* < 0.05 vs 0 μM GSK, One-Way ANOVA with Dunnett’s *post-hoc*; *n* = 15 zebrafish, 3 independent experiments. (**D**) Effect of muscle relaxant d-tubocurarine (dTC, 0.03–3 mM) on escape responses after head or tail touch at 76 hpf; Grey bands are mean ± SEM for control zebrafish; \**P* < 0.05 vs 0 mM dTC, One-Way ANOVA with Dunnett’s *post-hoc, n* = 15 zebrafish, 3 independent experiments. (**E**) (*i-iv*) Representative images of zebrafish for the indicated-treatments. (*v*) Quantification of zebrafish with morphological defects at 76 hpf for the indicated treatments. \**P* < 0.05 vs control, One-Way ANOVA with Sidak’s *post-hoc*, *n* = 15 zebrafish; 3 independent breedings. Blue arrows: curly-up tail.

### 3.3 PERK inhibition induces morphologic defects in zebrafish

PERK loss-of function elicits skeletal dysplasia in patients (5). To investigate the role of PERK in skeletal development, we exposed zebrafish embryos to the PERK inhibitor from 4 to 76 hpf and monitored their development over time. GSK (10 μM) evoked a significant time-dependent increase in morphological defects (Figure 3 A *i*). The most common defect was a curly-up tail and a few cases of scoliosis (Figures 3 A *ii* - *iv*). Exposure to GSK (10 μM) caused 20 ± 5 degrees change in the zebrafish trunk-tail angle at 76 hpf (Figure 3 B) and a significant decrease in body length (Figure 3 C).

To ascertain if the morphological defects evoked by GSK originate primarily in developing skeletal or muscular structures, we tested if the defects could be reversed by a muscle relaxant (d-tubocurarine; dTC; antagonist of muscular nicotinic receptors (39); effective in zebrafish (40)). We observed a concentration-dependent decrease in escape responses, with a maximal relaxation effect at 3 mM (Figure 3 D *i,ii*), confirming dTC efficacy as previously described (40). 3 mM dTC did not recover trunk-tail angle in GSK-treated zebrafish (Figure 3 E), indicating that that abnormal skeletal development, rather than muscle contraction, is the main cause of the morphological defects.

We measured otolith area (Supplementary Figure 1) and eye/body ratio (Supplementary Figure 2) as negative controls (unrelated with WRS phenotypes) and found no changes upon GSK exposure indicating that GSK effects on zebrafish morphology are selective.

These results indicate that upon PERK inhibition, zebrafish presented abnormal skeletal development and growth retardation, indicating that this small vertebrate holds potential to study the role of PERK in skeletal development.

### 3.4 PERK inhibition delays endocrine pancreas development and limits glucose uptake in zebrafish

Lack of PERK in WRS patients leads to decreased !-cell mass, impaired insulin production and neonatal diabetes (41, 42). Exocrine pancreas dysfunction was found in only 10.6% of patients (43, 44). Thus, we explored if zebrafish could be useful to study PERK’s role in pancreatic development.

To ascertain the effects of PERK inhibition in exocrine and endocrine pancreas development, we used two transgenic zebrafish lines expressing GFP in insulin-producing *β*-cells (Figure 4 A): one line also expressed mCherry in somatostatin-producing “-cells from the endocrine pancreas – Tg(*ins:GFP, sst:mCherry*); the other line expressed mCherry in elastase-producing exocrine pancreas –Tg(*ins:GFP, ela:mCherry*). We exposed both lines to GSK (10 µM) and measured the GFP/mCherry labelled areas as proxies for cell mass.

**Figure 4.**
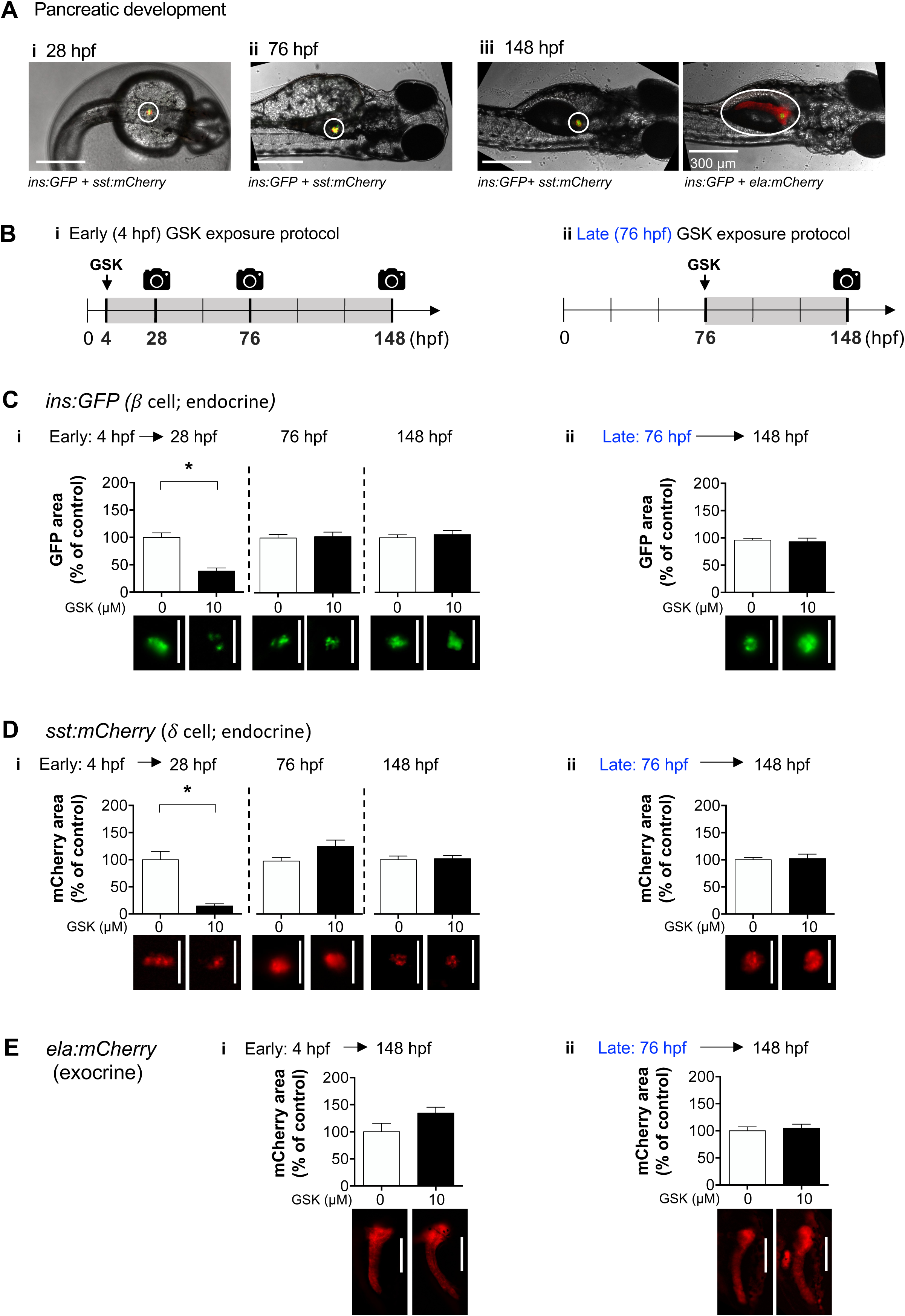
Pancreatic development in transgenic zebrafish. **(A)** Representative images of normal pancreas development in Tg(*ins:GFP, sst:mCherry*) and Tg(*ins:GFP, ela:mCherry*) zebrafish. (**B)** early (*i*) and late (*ii*) GSK2606414 (GSK)-exposure protocols. (**C-E**) Area measurements for *ins:GFP* (green, insulin-producing !-cells), *sst:mCherry* (red, somatostatin-producing “-cells), and *ela:mCherry* (red, elastase-producing exocrine cells), in zebrafish exposed to 10 μM GSK since 4 hpf (*i*, early exposure) or since 76 hpf (*ii*, late exposure). \**P* < 0.05, *t*-test, *n* = 4-33 larvae. Representative images of GFP and mCherry are shown below respective bars. Scale bars indicate 200 μm.

To test if the effects of GSK inhibition were selective for the early embryo development, we used two exposure protocols: the *early exposure protocol* (Figure 4 B *i*) started before the formation of the first Langerhans islet (endocrine pancreas); the *late exposure protocol* started after islet formation (Figure 4 B *ii*). Early PERK inhibition (4 hpf) delayed development of the endocrine pancreas, significantly decreasing *β* and “ cell mass at 28 hpf, which seems to recover overtime (Figure 4 C *i*, D *i*). Late inhibition (76 hpf) failed to alter endocrine cell mass (Figure 4 C *ii*, D *ii*). Regarding the exocrine pancreas mass, *ela:mCherry* fluorescence was only detectable at 148 hpf, and remained unchanged with either early or late

GSK exposure protocols (Figure 4 E). These findings support the hypothesis that PERK inhibition delays the early embryonic development of the endocrine pancreas.

To test if delayed endocrine development affected glucose homeostasis, we measured the uptake of the fluorescent glucose probe 2-NBDglucose (29) in zebrafish treated with GSK from 4 to 76 hpf (early exposure protocol). 2-NBDglucose uptake was measured in neuromast cells, given their direct exposure to probes in the external media, using Mitotracker Deep Red as a positive control for neuromast staining. Early PERK inhibition significantly decreased the uptake of 2-NBDglucose (Figure 5 A, B), indicating that PERK inhibition, induces a diabetic-like phenotype in zebrafish.

**Figure 5.**
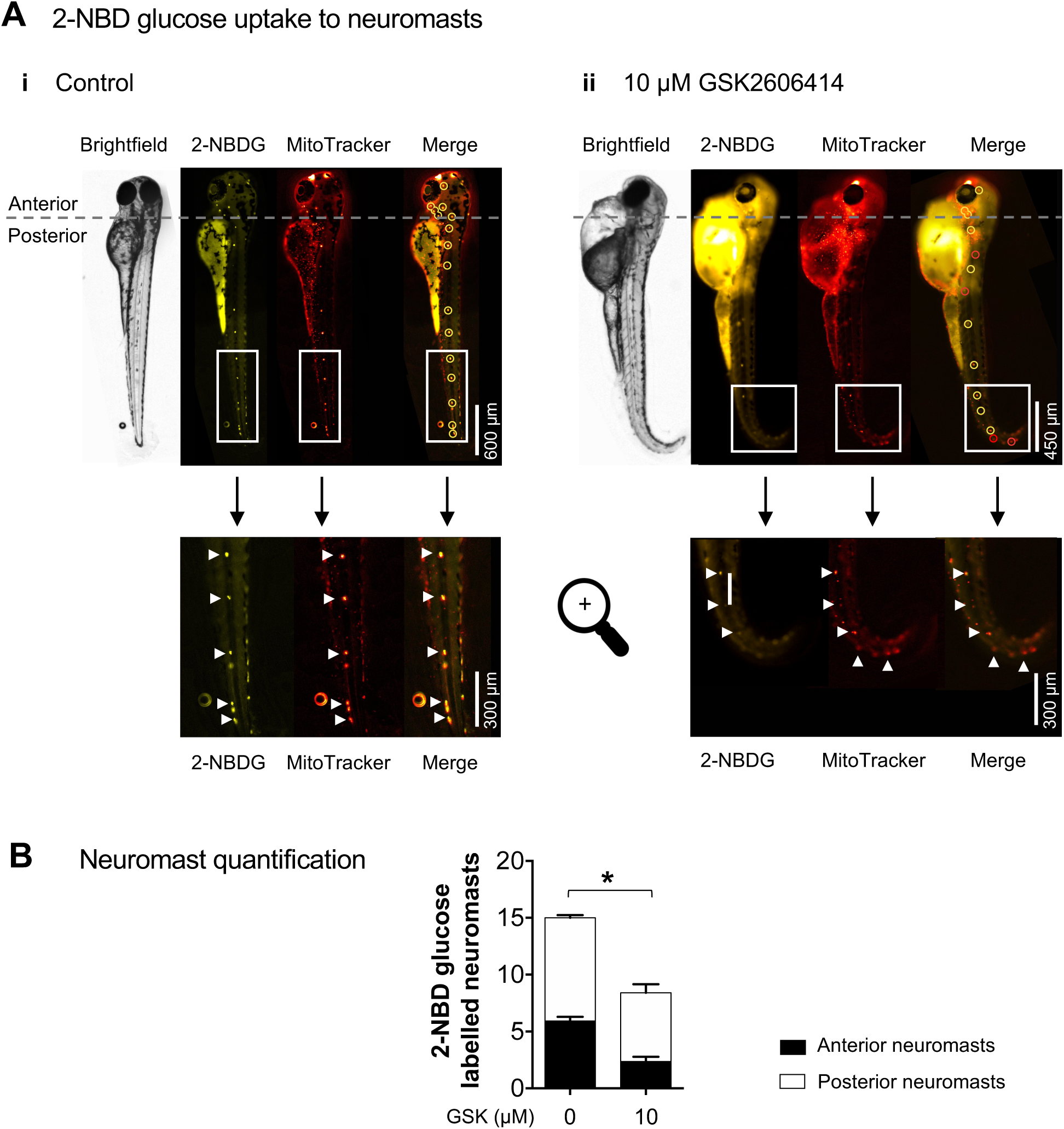
GSK2606414 effects on 2-NBDglucose uptake in zebrafish. (**A**) Representative brightfield and fluorescence images of zebrafish at 76 hpf, following treatment with (*i*) 0 μM (control) or (*ii*) 10 μM GSK2606414 since 4 hpf. Images show functional labelling at 76 hpf with the fluorescent glucose analogue 2-NBDglucose (2-NBDG) for neuromasts that uptake glucose, and with MitoTracker Deep Red that labels mitochondria in neuromasts as positive control. In the “Merge” images, yellow circles highlight neuromasts with both 2-NBDglucose and MitoTracker, while red circles highlight those containing MitoTracker only. Below, 2x amplified images show neuromasts from the zebrafish lateral line, with arrows indicating neuromasts on the side proximal to the microscope objective (the paired neuromasts on the opposite side are unmarked). (**B**) Quantification of neuromasts labelled with 2-NBDglucose. \**P* < 0.05, t-test, *n* = 9-12 zebrafish per condition.

### 3.5 PERK inhibition evokes neuromuscular and cardiac impairments in zebrafish

Beyond diabetes and skeletal dysplasia, patients with loss-of-function mutations in PERK may also present impaired brain development (45), muscle weakness (46) and cardiac alterations (47, 48). We first assessed the role of PERK on zebrafish neuromuscular function by measuring spontaneous tail-coiling and touch-evoked escape responses upon PERK inhibition (49–51). GSK (10 µM) significantly increased tail-coiling events in early development (28 hpf embryos inside the chorion; Figure 6 A) and significantly decreased escape responses in later development (76 hpf larvae; Figure 6 B). To test for altered muscle integrity, we assessed the arrangement of sarcomeres by measuring how well they diffracted polarized light (birefringence assay) (52) (Figure 6 C *i*). Treatment with 10 μM GSK significantly decreased muscle birefringence intensity (Figure 6 C *ii, iii*).

**Figure 6.**
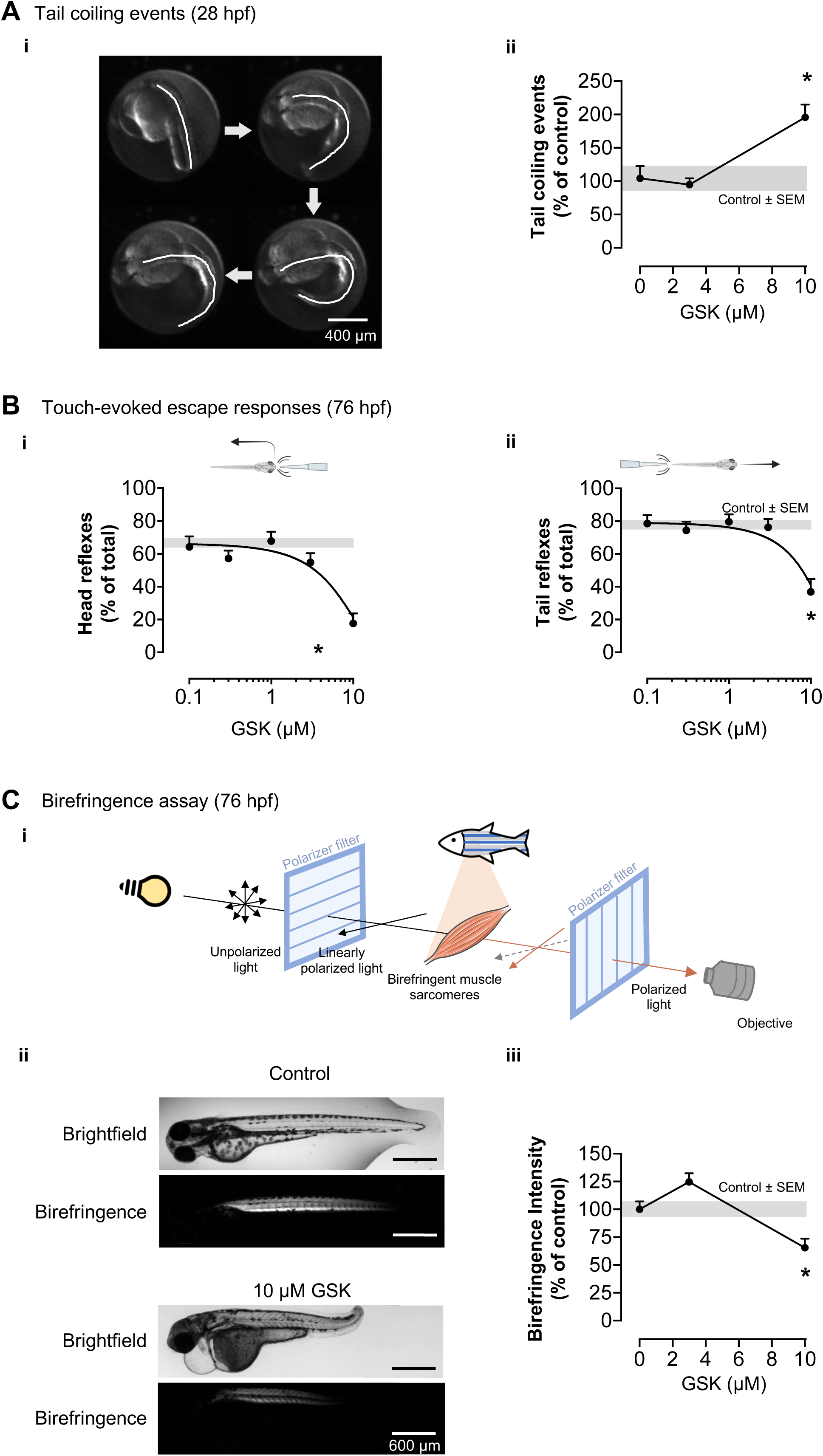
GSK2606414 effects on neuromotor responses and muscle integrity. (**A**) (*i*) Representative tail coiling time-lapse: white line highlights notochord movement. (*ii*) Quantification of tail coiling in 28 hpf zebrafish treated with GSK2606414 since 4 hpf; *n* = 23-26 zebrafish. (**B**) Quantification of escape responses after head (*i*) or tail (*ii*) touches at 76 hpf; *n* = 13-114 zebrafish. (**C**) (*i*) Schematic representation of the birefringence assay for muscle integrity; (*ii*) Representative images of muscle birefringence at 76 hpf. (*iii*) Quantification of birefringence intensity; *n* = 15 zebrafish. (**A-C**) Grey bands are mean ± SEM for 0 μM GSK (control); \**P* < 0.05 vs. (control), One-Way ANOVA with Dunnett’s *post-hoc*.

Next, we monitored cardiac morphology and function in PERK-inhibited zebrafish, where 10 μM GSK induced a time-dependent increase in pericardial oedema (Figure 7 A). Additionally, at 76 hpf, 10 μM GSK induced bradycardia without altering atrio-ventricular coordination in zebrafish (Figure 7 B).

**Figure 7.**
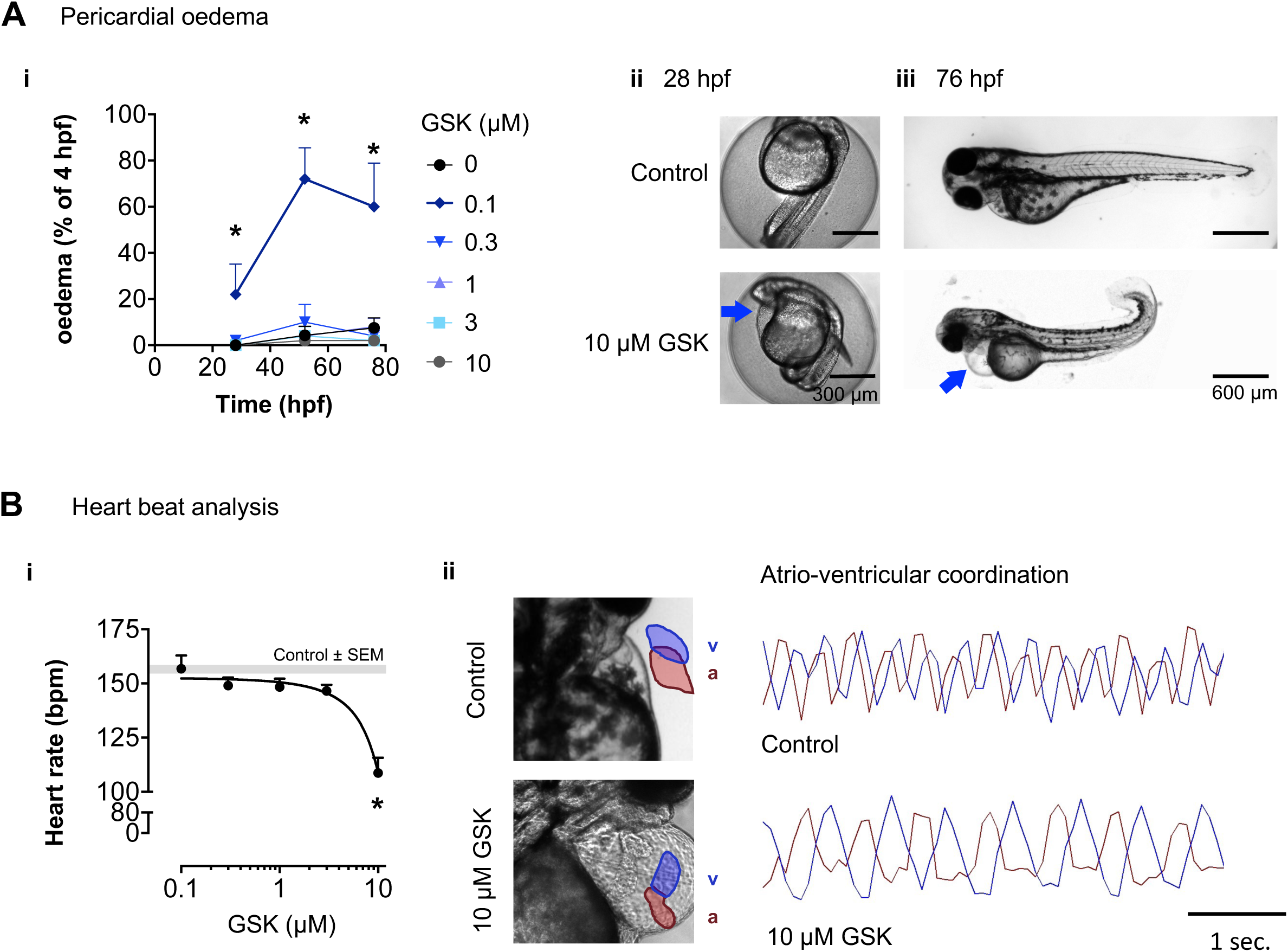
GSK2606414-induced morphological and functional cardiac defects. (**A**) (*i*) Quantification of live zebrafish with pericardial oedema at the indicated hpf and GSK concentrations. \**P <* 0.05, 10 vs. 0 µM GSK (control); Two-Way ANOVA with Sidak *post-hoc n*= 50-234 zebrafish; 5 independent experiments. (*ii,iii*) Representative images of zebrafish treated with GSK vs. control; blue arrow: pericardial oedema. (**B**) (*i*) Heartbeats per minute (bpm) of 76 hpf zebrafish exposed to increasing GSK concentrations since 4 hpf; Grey band is mean ± SEM for control; \**P* < 0.05 vs control, One-Way ANOVA with Dunnett’s *post-hoc*, *n* = 13-116 zebrafish; (*ii*) Representative images of the heart and plots of atrio-ventricular coordination in control and 10 μM GSK-treated zebrafish (a: atrium – red; v: ventricle – blue).

These results suggest that PERK-inhibited zebrafish present neuromuscular and cardiac abnormalities in morphology and function, indicating that zebrafish are suitable models to study PERK effects on neuromuscular and cardiac development and function.

## 4. Discussion

This study contributes to the understanding of PERK’s role in the embryonic development of zebrafish, building on PERK’s functions in zebrafish sleep and oxidative stress regulation (15–17). We show that PERK inhibition causes defective skeletal development and diabetic-like phenotypes, cardiac and neuromuscular alterations in zebrafish, mirroring key features of Wolcott-Rallison syndrome patients. These results highlight the potential of zebrafish to study PERK’s role in developmental diseases with multisystem involvement such as the WRS.

PERK inhibition with GSK2606414 (GSK), one of the most used PERK inhibitors in the scientific literature, decreased ISR activation in animals like mice (53), frogs (54) and flies (15). In the present study, we show that GSK decreases the levels of ISR markers in zebrafish, which suggests that GSK is bioactive and effective in decreasing PERK pathway activity. Additionally, GSK evoked key WRS-associated phenotypes (impaired skeletal development and diabetic-like phenotype), while sparing non-WRS-associated parameters as otolith area and eye/body ratio. Although we cannot exclude GSK off-targeting c-KIT and RIPK kinases (55, 56), zebrafish with genetically inhibited KITa/b and RIPK3 (57, 58) did not show the curly-up tail and other structural alterations reported in our study, suggesting that such alterations are not off-targets effects of GSK in c-KIT and RIPK kinases. Thus, these results are compatible with GSK selectively inhibiting PERK in zebrafish.

Loss of PERK function causes growth retardation and skeletal dysplasia in humans (5) and mice, via impairments in collagen secretion, osteoblast differentiation and bone mineralization (9, 59, 60). Here, we show that PERK inhibition in zebrafish causes growth retardation and spinal deformities resistant to muscle relaxation with tubocurarine. Since at the time of the assays (76 hpf) the zebrafish cartilage is starting to ossify (61), our findings indicate that PERK inhibition impairs zebrafish skeletal development.

PERK controls pancreatic !-cell development and insulin production (62). Loss of PERK function in humans and mice causes decreased pancreatic mass and neonatal diabetes, by impairing !-cell proliferation and proinsulin maturation in the endoplasmic reticulum (63–66). Here we show that PERK inhibition during early zebrafish development delays endocrine pancreas formation, resulting in a decreased !-cell mass, which gradually recovers overtime. To assesses whether glucose homeostasis is impaired in PERK-inhibited zebrafish we used the fluorescence glucose uptake probe 2-NBDglucose. We found reduced 2-NBDglucose uptake, which indicates that glucose homeostasis remains partly compromised despite the ongoing !- cell mass recovery. Such recovery in the presence of GSK suggests that zebrafish can regenerate pancreatic tissue (67, 68) by PERK-independent mechanisms. The identification of such PERK-independent mechanisms in zebrafish may assist the development of novel therapies for !-cell regeneration for human diabetes (69).

In contrast to the delayed development of the zebrafish endocrine pancreas, we found no effect of PERK-inhibition upon the mass of the exocrine pancreas. While different mechanisms regulate development of the endocrine and exocrine pancreas (43, 65), we cannot exclude that extended exposure of zebrafish to the PERK inhibitor may alter the mass of the exocrine pancreas. Future studies using selective PERK knockout in zebrafish pancreatic cell sub-types may contribute to understand differential pancreatic vulnerability.

Beyond pancreatic dysfunction, PERK inhibition in humans and mice impairs neuronal and muscular functions (5, 6, 11, 70, 71). Here we show that PERK inhibition in zebrafish increases spontaneous tail movements early in development, which arise in a spinal central pattern generator that provides rhythmic depolarizations (49, 51), and decreases touch-evoked escape responses later in development, which arise in sensory neurons (50, 51). We also show that PERK-inhibited zebrafish display altered muscular integrity, likely contributing to decreased escape responses. Thus, as with mammals, the mechanisms underlying motor impairment in PERK-inhibited zebrafish combine dysfunctions in neurons and muscle.

Patients and mice with PERK loss of function often display cardiac malformations (10, 47, 48). Here we show that PERK-inhibited zebrafish display cardiac abnormalities, such as oedema and bradycardia. The transparency and small size of young zebrafish is amenable to the detection of cardiac abnormalities that would be fatal in larger animals due to insufficient oxygen diffusion (72). Moreover, zebrafish may assist discovery of heart regeneration mechanisms (73). Thus, future studies with a more detailed structural and functional characterization of the cardiac abnormalities in PERK-inhibited or deleted zebrafish, may provide valuable insights into disease mechanisms and potential therapeutics.

## 5. Conclusions

This study highlights the potential of the small vertebrate zebrafish as a model for developmental diseases, namely those caused by PERK inhibition like the WRS. We show that pharmacological inhibition of PERK with GSK2606414 mimicked WRS pathophysiology by inducing defective growth and skeletal development, and diabetic-like phenotypes such as delayed development of the endocrine pancreas and altered glucose homeostasis. PERK inhibition also impaired neuromuscular and cardiac development, while sparing non-WRS associated parameters like otolith area and eye/body ratio. Future studies using CRISPR/Cas9 technology can be used to generate zebrafish models with conditional/selective expression of human PERK with WRS-driving mutations (74). Also, future research on the mechanisms governing PERK-independent pancreatic regeneration in zebrafish may uncover potential therapeutic targets for diabetes. Further, the scalability of zebrafish models holds great potential to enhance the efficiency of drug screening against PERK-associated developmental diseases.

## Declarations Authorship contribution

**Conceptualization** – LA, BRP, JMAO**; Methodology –** LA, NASO, JB, BRP, JMAO; **Investigation –** LA, LPL, NASO, RFOS, BS; **Validation** – LA, LPL, NASO, RFOS, BS; **Data Curation** – LA; **Formal Analysis** – LA, LPL, NASO, RFOS; **Visualization** – LA, LPS, NASO; **Writing – original draft** – LA; **Writing – review and editing** – LA, LPL, NASO, JB, BRP, JMAO; **Resources** – JB, JMAO; **Funding acquisition** – JB, JMAO; **Project administration** – JMAO; **Supervision –** BRP, JMAO.

## Data availability statement

The datasets generated during and/or analysed during the current study are available from the corresponding author upon reasonable request.

## Abbreviations List

2-NBDglucose: (2-(7-Nitro-2,1,3-benzoxadiazol-4-yl)-D-glucosamine)
ATF4: Activating Transcription Factor 4
CHOP: C/EBP homologous protein
dpf: days post-fertilization
dTC: d-tubocurarine
eIF2α: eukaryotic translation initiation factor 2α
ER: Endoplasmic reticulum
hfp: hours post-fertilization
GSK: GSK2606414
ISR: Integrated Stress Response
NPH: Nikon-Prior-Hamamatsu
PERK: Protein kinase R-like ER kinase
WRS: Wolcott-Rallison Syndrome

## Supplementary Material

**Supplementary Figure 1.**
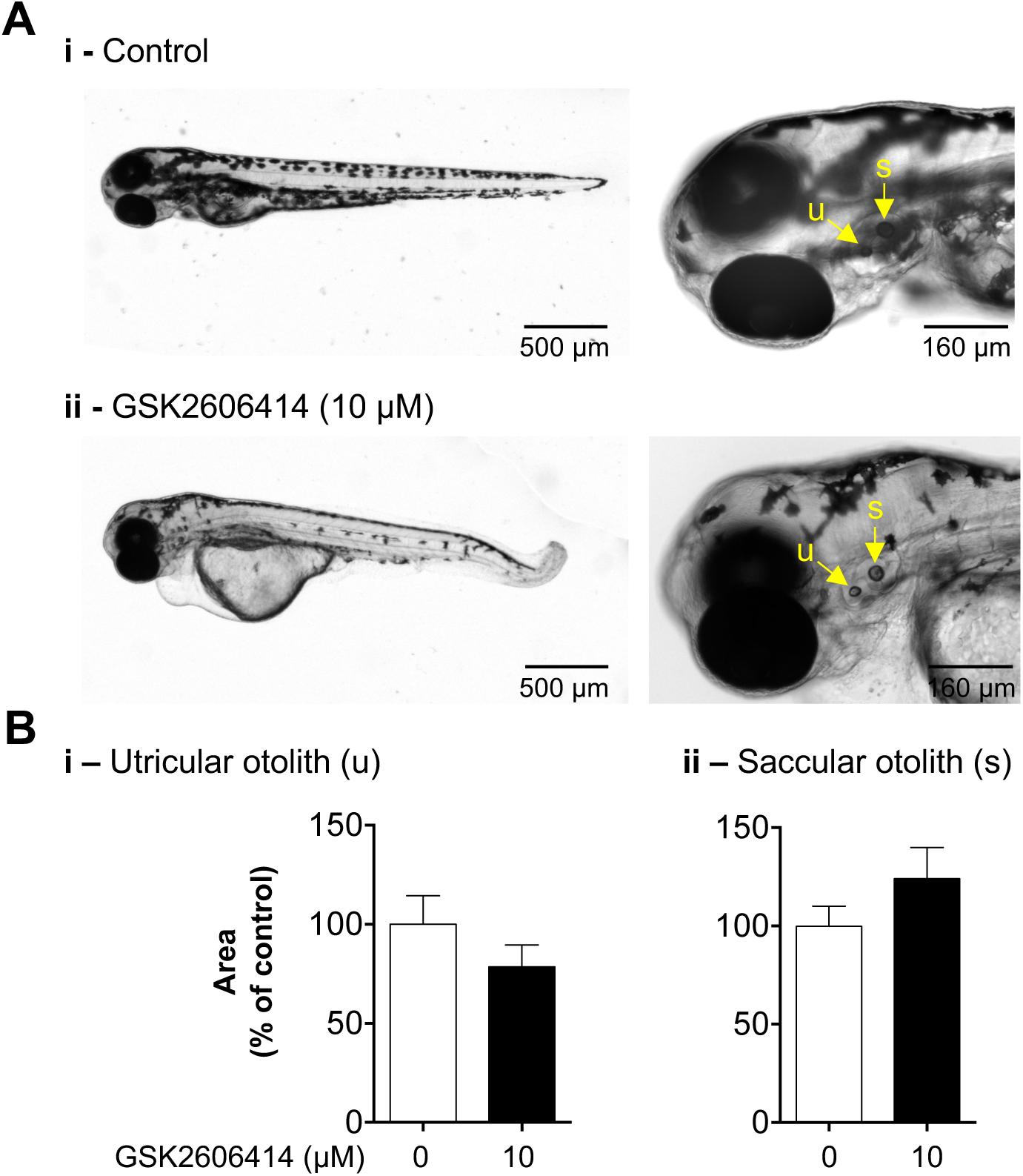
GSK2606414 (GSK) does not alter zebrafish otolith area. **(A)** Representative images of zebrafish at 76hpf following treatment with control (0 μM GSK) or 10 μM GSK since 4 hpf. Arrows: (u) utricular/anterior otolith, (s) saccular/posterior otolith. **(B)** Mean ± SEM of the otolith area, *P* > 0.05, t-test, *n* = 15 larvae, 3 zebrafish experiments.

**Supplementary Figure 2.**
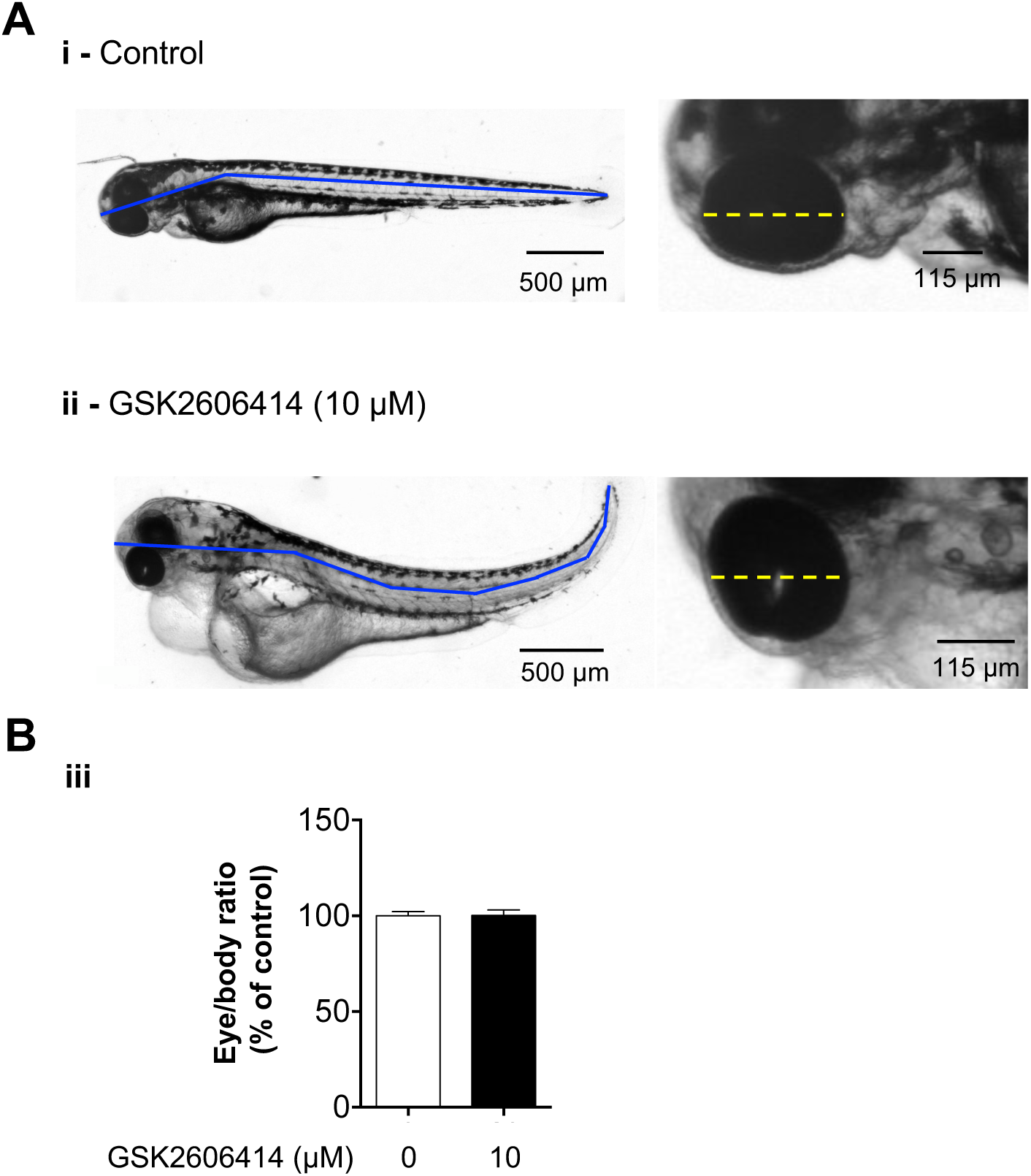
GSK2606414 does not alter zebrafish eye/body ratio. **(A)** Representative images of zebrafish at 76hpf following treatment with control (0 μM GSK) or 10 μM GSK since 4 hpf. Solid blue lines: body length; dashed yellow lines: eye diameter. **(B)** Mean ± SEM of the eye/body ratio, *P* > 0.05, t-test, *n* = 15 zebrafish, 3 independent experiments.

**Supplementary Table 1.**
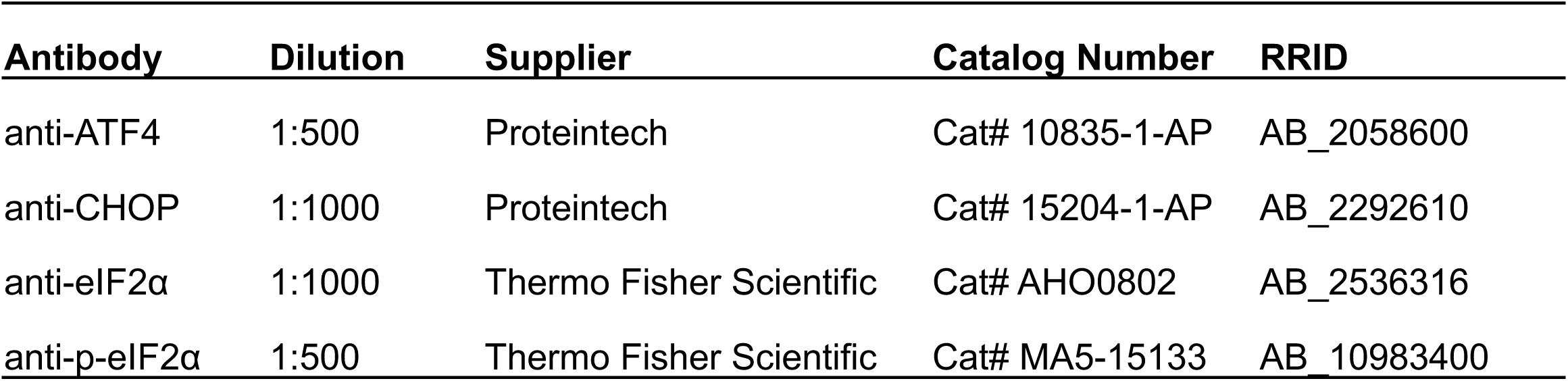
Primary antibodies used in western blots from Fig. 2 D.

| Antibody | Dilution | Supplier | Catalog Number | RRID |
| --- | --- | --- | --- | --- |
| anti-ATF4 | 1:500 | Proteintech | Cat# 10835-1-AP | AB_2058600 |
| anti-CHOP | 1:1000 | Proteintech | Cat# 15204-1-AP | AB_2292610 |
| anti-eIF2α | 1:1000 | Thermo Fisher Scientific | Cat# AHO0802 | AB_2536316 |
| anti-p-eIF2α | 1:500 | Thermo Fisher Scientific | Cat# MA5-15133 | AB_10983400 |

**Supplementary Table 2.** Secondary antibodies used in western blots from Fig. 2 D.

| Antibody | Dilution | Supplier | Catalog Number | RRID |
| --- | --- | --- | --- | --- |
| anti-mouse | 1:4000 | Thermo Fisher Scientific | Cat# G-21040 | AB_2536527 |
| anti-rabbit | 1:4000 | Thermo Fisher Scientific | Cat# G-21234 | AB_2536530 |

